# Physiological phenotypes differ among color morphs in introduced populations of the common wall lizard, *Podarcis muralis*

**DOI:** 10.1101/2023.06.17.545449

**Authors:** Ali Amer, Sierra Spears, Princeton L. Vaughn, Cece Colwell, Ethan H. Livingston, Wyatt McQueen, Anna Schill, Dustin Reichard, Eric J. Gangloff, Kinsey M. Brock

## Abstract

Many species exhibit color polymorphisms which have distinct physiological and behavioral characteristics. However, the consistency of morph trait covariation patterns across species, time, and ecological contexts remains unclear. This trait covariation is especially relevant in the context of invasion biology and urban adaptation. Specifically, physiological and behavioral traits pertaining to energy maintenance are crucial to fitness, given their immediate ties to individual reproduction, growth, and population establishment. We investigated the physiological and behavioral traits of *Podarcis muralis,* a versatile color polymorphic species that thrives in urban environments (including invasive populations in Ohio, USA). We measured five physiological and behavioral traits (circulating corticosterone and triglycerides, hematocrit, body condition, and field body temperature) which compose an integrated multivariate phenotype. We then tested variation among stable color polymorphisms in the context of establishment in an urban environment. We found that the traits describing physiological status and strategy shifted across the active season in a morph-dependent manner-the white and yellow morphs exhibited clearly different multivariate physiological phenotypes, characterized primarily by differences in circulating corticosterone. This suggests that morphs have different strategies in physiological regulation, the flexibility of which is crucial to urban adaptation. The white-yellow morph exhibited an intermediate phenotype, suggesting an intermediary energy maintenance strategy. Orange morphs also exhibited distinct phenotypes, but the low prevalence of this morph in our study populations precludes clear interpretation. Our work provides insight into how differences among stable polymorphisms exist across axes of the phenotype and how this variation may aid in establishment within novel environments.

**Epigraph:** “*If you are only moved by color relationships, you are missing the point*.”
– Mark Rothko, Latvian-American abstract painter

## Introduction

Biological diversity is threatened by habitat fragmentation, large-scale human landscape modification, and environmental degradation (Liu *et al*. 2016). However, populations that are more phenotypically diverse should be less vulnerable to environmental change and genetic bottlenecks (Forsman & Åberg 2008; Forsman *et al*. 2008). Color polymorphism is a form of extreme intraspecific diversity, where two or more genetically-determined color morphs coexist within a single population. Though these color morphs occur in strict syntopy, interact with each other, and interbreed (Pérez i de Lanuza & Carretero 2018), many morphs evolve so-called ‘alternative phenotypes’ that comprise multiple traits. Alternative color morph phenotypes consist of multivariate differences in traits such as behavior (Abalos *et al*. 2016; Brock *et al*. 2022b; Sinervo & Lively 1996; Yewers *et al*. 2016), morphology (Huyghe *et al*. 2007), ecology (Lattanzio & Miles 2014), physiology (Comendant *et al*. 2003; Forsman 2000; Mills *et al*. 2008), and performance (Huyghe *et al*. 2009; Zajitschek *et al*. 2012), which may influence fitness (Stuart-Fox *et al*. 2021). Morph-specific phenotypic differences may promote the use of diverse environmental resources that aid in the successful establishment in novel environments and long-term persistence of color polymorphic species through reduced intraspecific competition (Forsman 2016; Forsman *et al*. 2008; Forsman & Wennersten 2016; Pizzatto & Dubey 2012; Svensson 2017; but see *Bolton et al.* 2016). Thus, polymorphic species may be primed for range expansion (Forsman *et al*. 2008), and characterizing the phenotypic complexes associated with intraspecific color morphs is crucial to understanding how biological diversity is maintained and aids in population establishment.

Lizards of the genus *Podarcis* have been introduced and successfully established in various locations around the globe (Hedeen 1984; Heym *et al*. 2013; Michaelides *et al*. 2015; Podnar *et al*. 2005; Ribeiro & Sá-Sousa 2018; Silva-Rocha *et al*. 2014). In addition to these repeated invasions, almost all species in the genus exhibit high levels of color polymorphism (Brock *et al*. 2022a), making *Podarcis* an ideal study system to understand the role of discrete color polymorphisms in the context of invasion. Our study species, the common wall lizard (*Podarcis muralis*) has expanded beyond its native range of southern Europe (Speybroeck *et al*. 2016) and successfully established in many places, including Germany (Heym *et al*. 2013), England (Michaelides *et al*. 2015) and Cincinnati, Ohio, USA (Brown *et al*. 1995b; Davis *et al*. 2021; Deichsel & Gist 2001; Hedeen 1984). *P. muralis* were introduced to Cincinnati in the early 1950s, when, following a vacation to Northern Italy, a young boy released approximately 10 individuals into his yard (Brown *et al*. 1995b; Davis *et al*. 2021; Deichsel & Gist 2001; Hedeen 1984). Since then, they have exploded in population size, numbering in the hundreds of thousands of individuals across Cincinnati and the surrounding areas (J. Davis pers. comm). Additionally, Cincinnati is heavily urbanized, which alters the environment along many dimensions, including structural habitat (Mohan *et al*. 2011) and temperature (Chow & Roth 2006). These environmental alterations have large and wide-reaching effects on the populations that experience them, including influencing physiology (Bonier 2023; Campbell-Staton *et al*. 2020; Hall & Warner 2018; Isaksson 2020), morphology (Putman & Tippie 2020; Vaughn *et al*. 2021; Winchell *et al*. 2018), and behavior (Putman *et al*. 2020; Sparkman *et al*. 2018; Stroud *et al*. 2019). By researching these urban areas, scientists can gain insights into the various mechanisms underlying the persistence – or extirpation – of stable polymorphisms in a novel ecological context.

Species introduced into a novel environment often make a number of ecological and physiological adjustments in order to adapt to its new niche. One of the most important problems faced by ectotherms in urban environments is maintaining body temperature (Campbell-Staton *et al*. 2020; Diamond *et al*. 2017; Diamond *et al*. 2018; Sándor *et al*. 2021). Body temperature affects practically all aspects of an ecotherms’s life, including metabolic rate, locomotion, foraging, defending territory, and reproduction (Angilletta *et al*. 2002; Flouris & Piantoni 2015; Huey & Stevenson 1979; Navas & Bevier 2001). In order to maintain their preferred body temperature, ectotherms (like lizards) must select their environment carefully. To avoid intraspecific competition within an environment, lizards specialize and adapt to specific microhabitats (Badillo-Saldaña *et al*. 2022; Stuart-Fox *et al*. 2021). This specialization can be seen in species like *Podarcis erhardii,* where different color morphs are found in different microhabitats (BeVier *et al*. 2022). These microhabitats are thermally different, and the lizards that inhabit them must adjust their active body temperatures accordingly. Orange *P. erhardii* morphs for example, are more often found in cooler, shadier environments, and as such, prefer cooler body temperatures compared to white and yellow morphs (Thompson *et al*. 2023). This observation suggests that differences in morph preferred body temperatures can provide an element of intraspecific thermal flexibility, aiding their ability to adapt to novel environments (Huey *et al*. 2012; Kearney *et al*. 2009; Litmer & Murray 2019; Nowakowski *et al*. 2018).

A key aspect of success in urban environments for many organisms is endocrine flexibility, though we lack clear patterns of what makes for a successful endocrine profile in an urban environment (Bonier 2023). Faced with increased anthropogenic disturbances during urban expansion, organisms can adjust their physiology through coordinated hormonal and behavioral responses to maintain homeostasis. Often the first aspect researchers examine is the adrenocortical response through the hypothalamus-pituitary-adrenal (HPA) axis (Gangloff & Greenberg 2023; Landys *et al*. 2006; Sapolsky *et al*. 2000; Wingfield *et al*. 1998). Corticosterone (CORT), the primary glucocorticoid (GC) in ectothermic vertebrates, plays a multi-faceted role in organismal function by regulating resource allocation, energy metabolism, and recovery from acute and chronic stressors (MacDougall-Shackleton *et al*. 2019; Wingfield *et al*. 1998). Acute GC secretion is considered beneficial as it enables organisms to cope with the novel environmental challenges attributed to urban living by promoting self-maintenance behaviors and a heightened physiological state favoring immediate survival (Landys *et al*. 2006). In this heightened state, GCs promote the mobilization of energy stores via the breakdown of stored triglycerides to free fatty acids, resulting in a decrease in circulating triglycerides (Remage-Healey & Romero 2001). Triglycerides (TRIG), the most energy-dense of all macromolecules, provide a measure of energy availability and gluconeogenesis and thus provide a useful complement to measures of CORT (Neuman-Lee *et al*. 2015; Price 2017; Sykes & Klukowski 2009), but are seldom measured in a natural context (but see Blair *et al*. 2000). Urban environments can also present altered water availability, an additional stressor for organisms inhabiting this space. Hematocrit (Hct), a measure of relative volume of red blood cells to total blood, can provide a measure of hydration status (Moeller *et al*. 2017; Peterson 2002), though hematocrit can be affected by other factors as well (Bodensteiner *et al*. 2021b; Puerta *et al*. 1996). Nonetheless, interindividual variation in Hct may provide useful insight, especially in concert with other measures, of the overall health of animal populations in novel environments. Measures of body condition inferred from morphological measurements are commonly invoked as a proxy for energetic status and, by extension fitness, and are a valuable tool in estimating the health and physiological state of animal populations (Stevenson & Woods 2006; Warner *et al*. 2016; Weatherhead & Brown 1996). While variation in relative body mass may be attributable to adipose tissue, muscle, or water, body condition can provide useful links to health status and fitness (Brischoux *et al*. 2016; Donihue *et al*. 2022; Gangloff *et al*. 2019; Le Galliard *et al*. 2004). Identifying patterns of covariation in these physiological indicators – and how these patterns may differ among the discrete color polymorphs – can provide clues as to the functional significance of covariation in these traits.

The Common wall lizard (*Podarcis muralis*) exhibits three genetically-determined monochromatic color morphs (orange, yellow, and white; see Fig. 1) and their intermediates (orange-white, white-yellow, and yellow-orange) that exhibit myriad morphological, behavioral, and performance differences in its native range (Abalos *et al*. 2016; Andrade *et al*. 2019; Zajitschek *et al*. 2012). Though studied extensively, the adaptive or functional significance of color polymorphism in *Podarcis* lizards remains elusive (Abalos *et al*. 2020; Huyghe *et al*. 2010; Sacchi *et al*. 2015a). Multiple lines of evidence suggest that these color badges on the throat serve a signaling function (Brock *et al*. 2020; Brock *et al*. 2022b; Pellitteri-Rosa *et al*. 2014); however, because metabolic costs of synthesis vary among different pigments and trade-off differently with other physiological functions, the extent to which color signals individual quality is uncertain (Abalos *et al*. 2020). Previous work has found that orange morphs in *P. muralis* are larger and have lower immune function, endurance, and survival compared to white morphs (Calsbeek *et al*. 2010). Further, there is conflicting data describing how the different color morphs will respond to seasonal changes with white and yellow morphs exhibiting distinct yet inverse patterns of aggression and testosterone levels across contexts in the breeding season (Coladonato *et al*. 2020; Sacchi *et al*. 2017). Previous work has shown the breeding season to be the catalyst of the differential immune responses among *P. muralis* color morphs (Galeotti *et al*. 2010; Sacchi *et al*. 2007b). This discrepancy in the literature further makes for an excellent opportunity to study variation in morph-linked traits across the active season in *P. muralis*.

**Figure 1.**
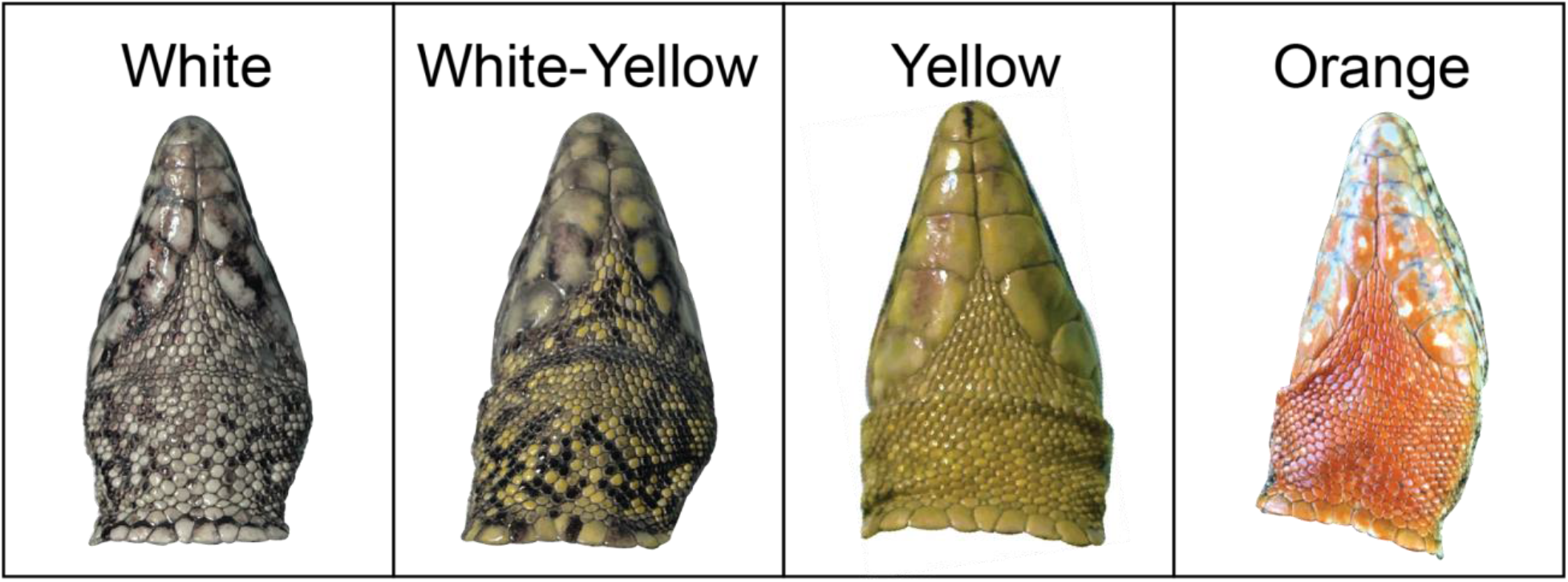
*Podarcis muralis* ventral color morphs as found in Ohio, USA.

Here, we investigated potential differences in physiological strategy among color morphs by measuring five physiological traits and behaviors related to energy balance and habitat selection – circulating corticosterone concentration (CORT), circulating triglyceride concentration (TRIG), blood hematocrit (Hct), selected body temperature (Tb), and body condition (mass relative to size). Our primary hypothesis is that wall lizards of different color morphs will differ in their physiological phenotype, thus suggesting that color morphs do not simply differ in appearance but in a variety of traits representing different strategies. From this follows multiple specific predictions: First, we predict the orange *P. muralis* morph in Cincinnati will be found in cooler and wetter habitats (as in orange morphs of other *Podarcis* species: BeVier *et al*. 2022; Pérez i de Lanuza & Carretero 2018; Thompson *et al*. 2023) and therefore will exhibit lower hematocrit and field body temperatures than the other morphs. Second, we predict that the white-yellow morph will serve as an intermediate physiological phenotype for the white and yellow morphs, as intermediate morphs in other studies (Brock *et al*. 2020; Brock *et al*. 2022b). Finally, we characterize shifts in the multivariate physiological-behavior phenotype of the color morphs in response to seasonal shifts in available temperatures, water, and food, and across the reproductive season. By investigating the potential differences in physiological traits and strategies employed by *P. muralis*, this paper is the first to examine and provide insight into how color morphs differ between key aspects of their physiology and ecology, as well as how these differences contribute to the success of adapting to a novel ecological environment.

## Materials and Methods

### Field data & lizard collection

We caught wall lizards (*Podarcis muralis,* Laurenti 1768) at six sites in Cincinnati, Ohio, USA and one site in Columbus, OH, USA during their peak activity and reproductive periods (08h00– 17h30; May 2020 – September 2020, June 2021 – October 2021; See Table S1 for complete sampling details). Air temperature (5 cm off the ground in the shade; PTH8708 Digital Temperature & Humidity Pen, General Tools, New York, USA) was collected at the beginning and end of each site survey. Lizards were captured via a thread lasso attached to an extendable fishing rod or by hand. We measured field body temperature (T_b_) by inserting a type K thermocouple approximately 0.5 cm into the lizard’s cloaca immediately after capture (< 10 s; HH801, Omega Engineering, Norwalk, Connecticut, USA). While field body temperatures are limited by thermoregulatory opportunities, we surveyed lizards during periods when conditions were optimal for thermoregulation, with low cloud cover and favorable air temperature (mean ± SD: 28.5 ± 3.24°C). We measured snout-vent length (SVL) as the distance from the tip of the snout to the posterior end of the anal scale with digital calipers (Model CD-6, Mitutoyo, Japan; mean ± sd: 61.8 ± 5.84, range: 46.74–74.21 mm). Lizards were weighed to the nearest 0.01 g using a digital scale (Weigh Gram Top-100, Pocket Scale, Tulelake, California, USA; mean ± sd: 5.9 ± 1.67, range: 2.50–9.34 g). One author (EJG) categorized each individual as orange, yellow, white, or white-yellow (Fig. 1). The color morphs are readily discernible by eye, based on throat and ventral scale colors (Calsbeek *et al*. 2010; Thompson *et al*. 2023). Ventral color polymorphism in *P. muralis* is discrete and consistent over time (Calsbeek *et al*. 2010; *Sacchi et al.* 2007a; Sacchi *et al*. 2007b). All research was conducted under Ohio Division of Wildlife Wild Animal Permit (23-014) and all procedures were approved by Ohio Wesleyan University IACUC (12-2020-02).

### Blood collection & processing

We collected a blood sample (20–45 μl) from the retro-orbital sinus (MacLean *et al*. 1973) using two heparinized glass capillary tubes per individual within < 4 minutes of capture (mean bleed time ± sd: 109.5 ± 46.8 s). We stored blood samples in capillary tubes on ice until processing. We spun the first capillary tube at 5000 *g* for 5 min on a centrifuge. We then measured the volume of packed red blood cells and total blood volume with digital calipers (Model CD-6, Mitutoyo, Japan). Hematocrit (Hct) was calculated as the ratio of packed red blood cells to total blood volume. We ejected the whole blood sample from the second capillary tube into microcentrifuge tubes, which we then spun at 3000 *g* for 5 min to separate plasma from red blood cells. We pipetted off the plasma, ejected it into fresh tubes, and flash-froze these tubes in liquid nitrogen. Plasma was then stored at −20°C until assays were performed (see below). We note that a subset of the field body temperature and hematocrit data were analyzed to address different questions in another manuscript (Spears *et al*. 2023, in review).

As our measure of body condition, we used the scaled mass index (SMI) as described by Peig & Green (2009), which accurately accounts for the allometric scaling of growth and has the advantage of producing an index in the same units as the measured mass (Brodeur *et al*. 2020). We first quantified the scaling exponent for our studied species *b_SMA_* by fitting an standardized major axis slope regression to log_10_-transformed data in accordance with the linearized power equation:

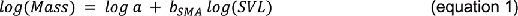

where *b_SMA_* is the slope. We expressed the scaled mass index of body condition (M) as follows

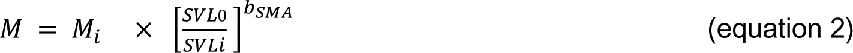

where M_i_ and SVL_i_ represent the individual body mass and snout-vent length respectively and SVL_0_ is the arithmetic mean snout-vent length of our study population (Peig & Green 2009).

### Laboratory Assays

Total plasma triglyceride concentration, defined as circulating levels of both triglycerides and free glycerols, was measured with a colorimetric assay (Triglyceride GPO Liquid Reagent Set, Catalog #23-666-410, MedTest Dx, Canton, Michigan, USA). Samples were run in duplicate at 1:2 dilution. Samples were re-run when the coefficient of variation (CV) for duplicate samples was above 15%, resulting in a mean sample CV of 4.56%. To assess inter-assay variability, we ran a pooled plasma sample in duplicate on each plate, providing a CV of 15.32%.

Plasma corticosterone (CORT) concentration was measured with a high sensitivity immunoassay (Corticosterone High Sensitivity EIA Kits, Immunodiagnostic Systems Inc., Scottsdale, Arizona, USA). Prior to CORT quantification, we validated the immunoassay by demonstrating parallelism of pooled wall lizard plasma sample dilution curves to a CORT control sample (test for heterogeneity of slopes: F_1,6_ = 0.642, P = 0.454). Plasma samples were run in duplicate at a 1:25 dilution. As with triglycerides, samples were re-run when the CV for duplicates was above 15%, resulting in a mean sample CV of 3.44%. We ran kit-provided controls in duplicate on each plate, though we were unable to run a common control or pool across all plates because plates were run by two different researchers (WM & AA) in different years. To test for potential researcher effects, we calculated the proportion of residual variance attributable to the researcher with a mixed linear model that included the researcher as a random effect and found this effect to be minimal (<0.00001%).

### Statistics: Multivariate Analyses

We analyzed the five physiological traits in a unified framework that allows us to simultaneously quantify the within-individual correlations of traits and test for differences in traits among color morphs and between sexes. TRIG and CORT were log_10_-transformed before analysis to meet the assumption of normal distribution of model residuals. We utilized a nonparametric multivariate analysis of variance (NP-MANOVA) with residual randomization in permutation procedure (RRPP) (Collyer & Adams 2018; Collyer & Adams 2022), and following Telemeco & Gangloff (2020). We created a model that included the categorical factors of color morph and sex, with significance determined from 999 iterations of the residual randomization procedure. Our initial model included the morph × sex interaction term, but we removed this because it was not significant (*P* = 0.469). We then extracted least-squares means in multidimensional space to compare physiological phenotypes among color morphs, conducted tests for differences in all pairwise color morph combinations, and extracted least-squares means and 95% confidence intervals for each of the traits included in the multivariate response matrix. We then tested for changes in physiological phenotypes across seasons using linear models, with PC scores from the first two axes of variation as dependent variables in separate models. We included the fixed effects of color morph (categorical factor with four levels: orange, white, yellow, and white-yellow), sex (categorical factor with two levels: male, female), sampling year (2020 or 2021), as well as the linear and quadratic effects of day of year (day since 1 January) and time of day (seconds past midnight). Initial models also included the interaction of color morph with the linear and quadratic effects of day of the year and the linear and quadratic effects of time of day. We performed backward selection, sequentially removing non-significant (*P* > 0.05) terms, beginning with the highest-order interaction terms, and then re-running the model. We assessed distributions of model residuals visually and with a Shapiro-Wilks test and determined the relative importance of fixed effects using type III sums of squares.

### Statistics: Univariate Analyses

We utilized linear models to test the influences of various factors on each of our five physiological variables: CORT, TRIG, Hct, T_b_ and SMI. As in the multivariate analysis, CORT and TRIG were log_10_-transformed before analysis. Model structure, backward selection procedure, and assessment of residuals was the same as above. All models met the assumption of normal distribution of residuals except for that of T_b_ (Shapiro-Wilks, P = 0.002), though visual inspection indicated only a slight skew which will minimally, if at all, affect parameter estimates and interpretation (Schielzeth *et al*. 2020). When the main effect of color morph significantly influenced the dependent variable, we conducted post-hoc comparisons of least-squares means with the emmeans package (Lenth *et al*. 2019). We conducted all statistical analyses in the programming language R (R Core Team 2023) with data figures created with ggplot2 (Wickham *et al*. 2023).

## Results

We collected a complete suite of phenotype data, including plasma corticosterone concentration (CORT), plasma triglyceride concentration (TRIG), hematocrit (Hct), field body temperature (T_b_), and standardized mass index (SMI), on a total of 86 lizards across the activity season from sites in Ohio, USA (**Supplemental Table 1**). Color morphs were not caught differently across day of the year (linear model, F_3,82_ = 1.06, *P* = 0.37) or time of day (F_3,82_ = 0.99, *P* = 0.40).

Our multivariate models (NP-MANOVA with RPPP) indicate that there is clear separation among color morphs (F_3,81_ = 2.00, *P* = 0.016) and between sexes (F_1,81_ = 4.30, *P* = 0.003) in the multivariate phenotype. Pairwise comparisons indicate significant differences between white and yellow morphs (*P* = 0.0008) with a trend for differences between orange and yellow morphs (*P* = 0.08; Table 1). The first two axes of variation among color morphs account for more than 96% of total variation between groups (Table 2, Fig. 2). PC1, accounting for 75.4% of variation, describes a continuum of lizards with high levels of CORT, TRIG, Hct, and SMI and a low T_b_, in contrast to lizards with the opposite combination of traits. PC2 accounts for 20.8% of total variation and contrasts lizards with high values of CORT, a high T_b_, and low TRIG, Hct, and SMI with lizards with the opposite combination of traits (Table 2, Fig. 2). Comparisons of least-squares means for each trait are presented in Fig. 3 and mean values for each trait by color morph are presented in Table 3.

**Figure 2.**
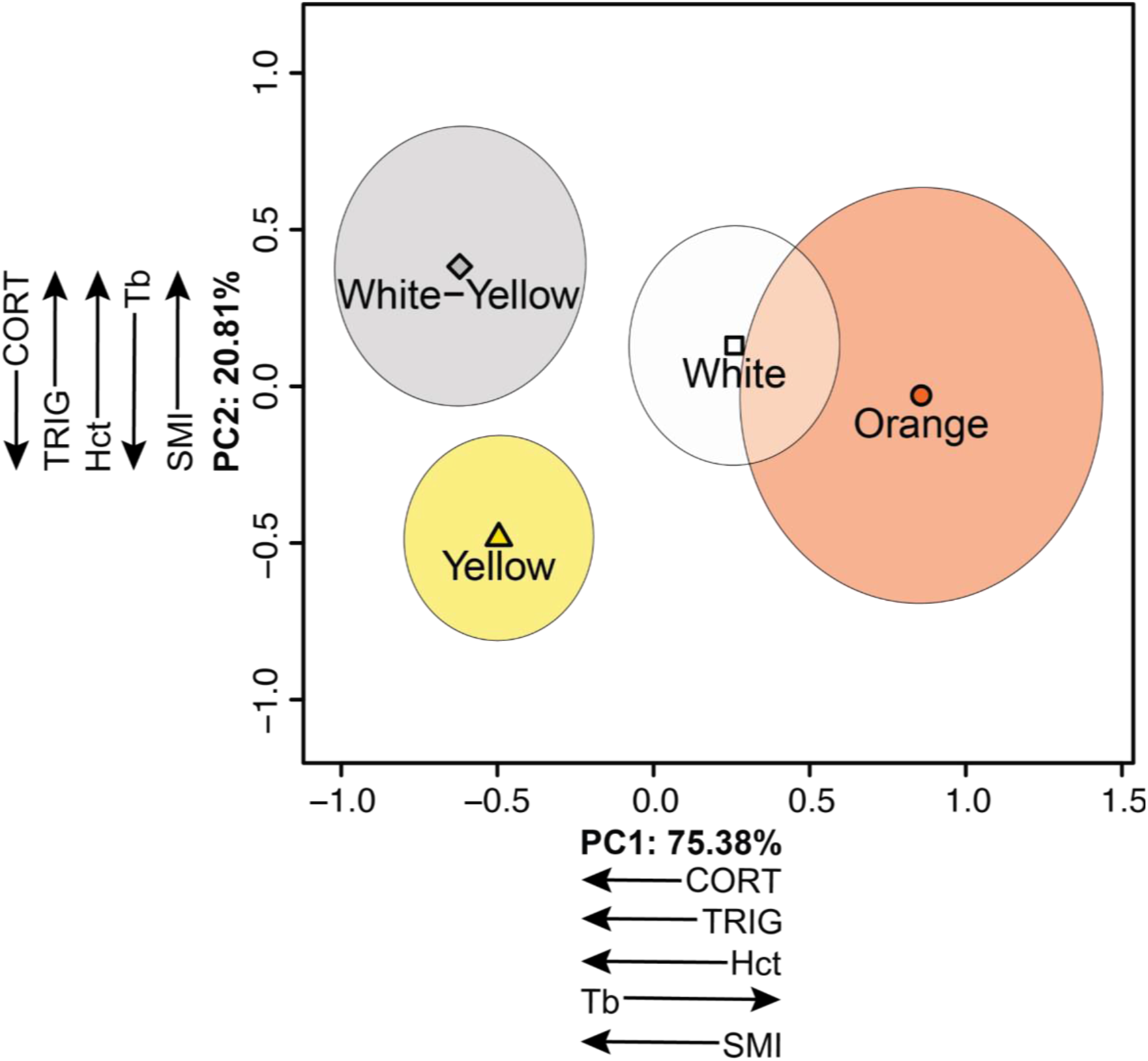
Principal component (PC) plots of phenotype of common wall lizards (*Podarcis muralis*) from Ohio, USA by color morph. Least-squares means and 95% confidence ellipses from non-parametric multivariate analysis of variance (NP-MANOVA) with randomized residuals in a permutation procedure (RRPP), including the fixed effects of color morph and sex (see main text for statistical details). Directionality of loadings for each PC axis are shown; full details provided in Table 2. Abbreviations: CORT = Plasma corticosterone concentration; TRIG = Plasma triglyceride concentration; Hct = Hematocrit; T_b_ = Field body temperature; SMI = Standardized mass index.

**Figure 3.**
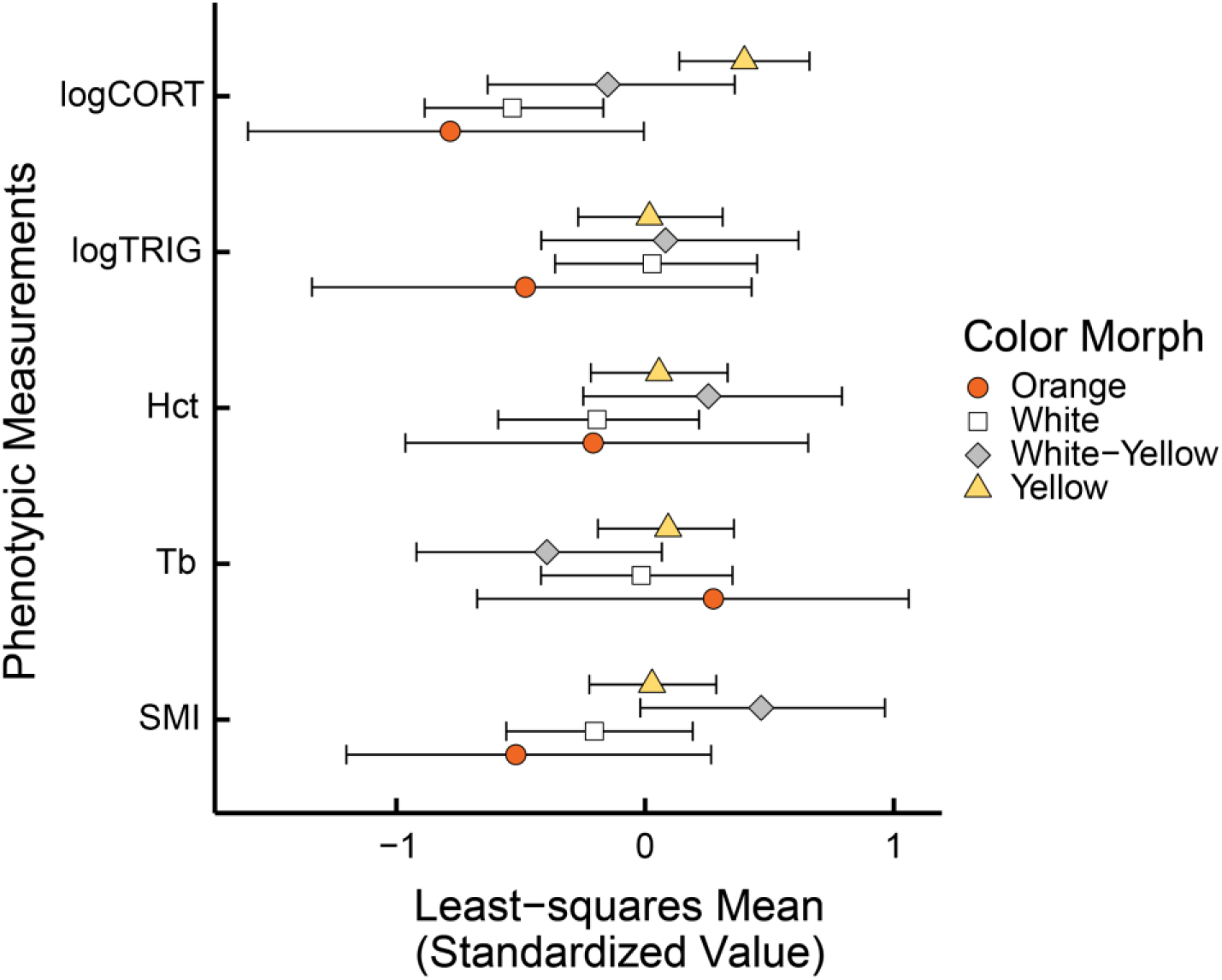
Least-squares means and 95% confidence intervals for phenotypic traits of common wall lizards (*Podarcis muralis*) from Ohio, USA generated from a non-parametric multivariate analysis of variance (NP-MANOVA) with randomized residuals in a permutation procedure (RRPP), including the fixed effects of color morph and sex (see main text for statistical details). Values shown are predicted from the model after accounting for covariation within the response matrix, displayed on a z-standardized scale. Abbreviations: CORT = Plasma corticosterone concentration; TRIG = Plasma triglyceride concentration; Hct = Hematocrit; T_b_ = Field body temperature; SMI = Standardized mass index.

**Table 1.**
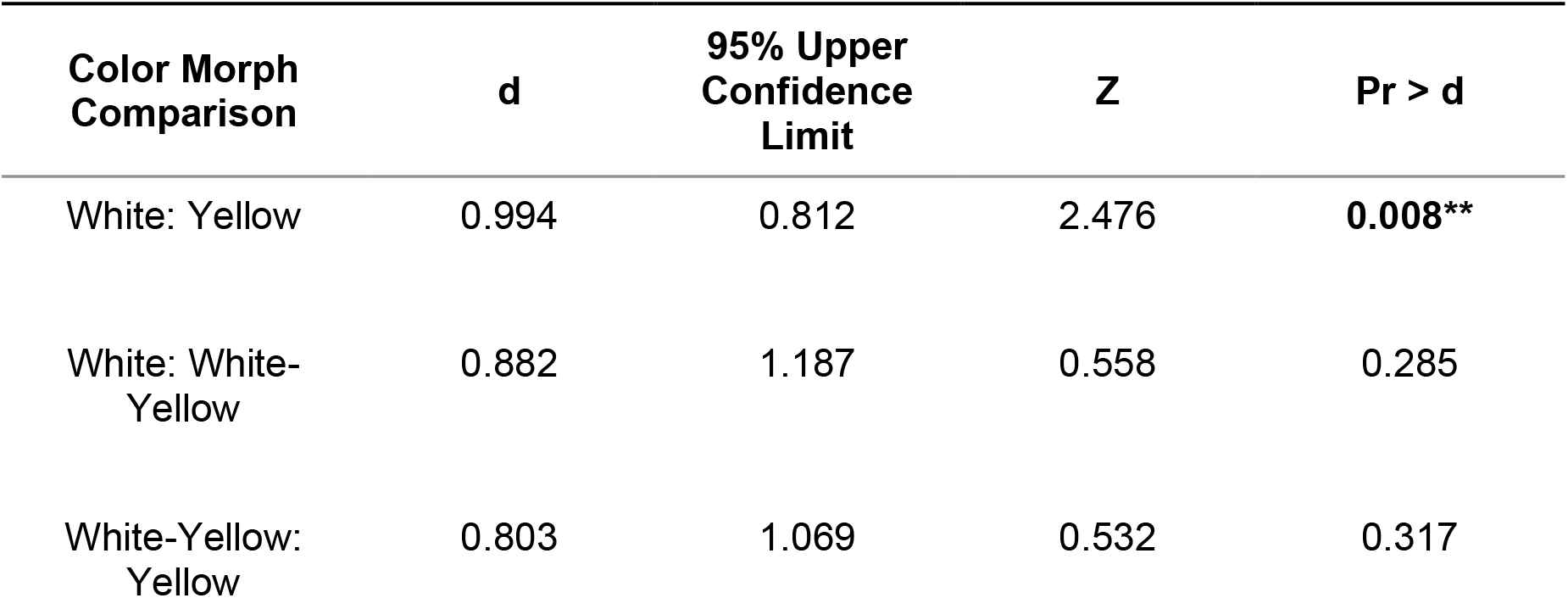
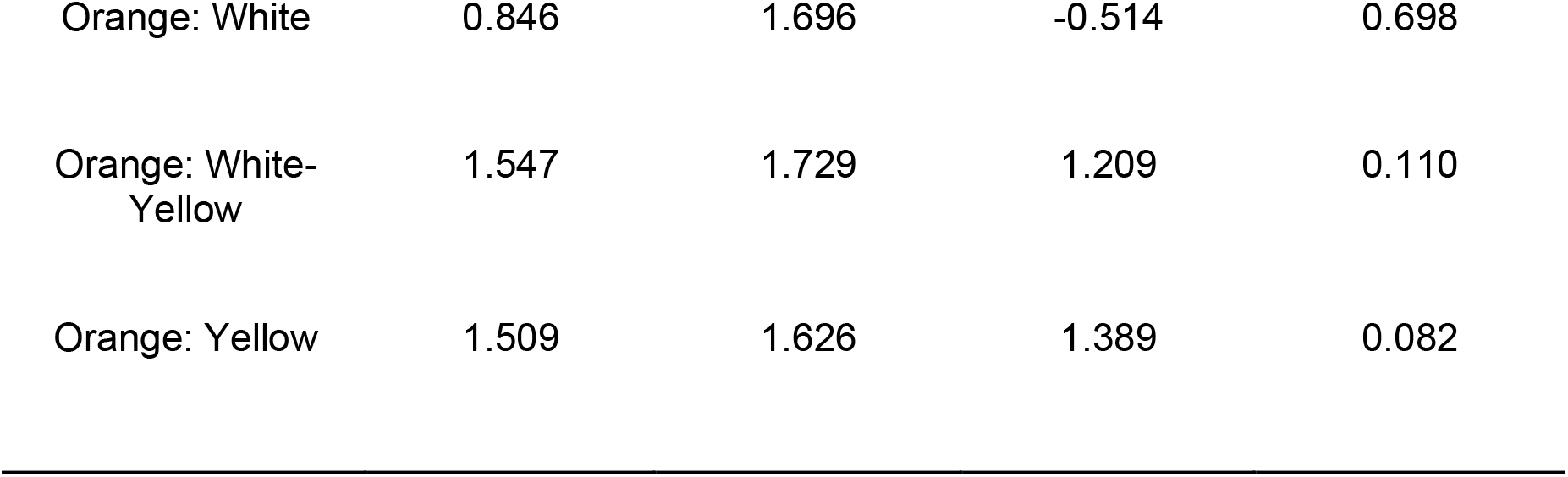
Pairwise comparisons of estimated least-squares means among color morph combinations from non-parametric multivariate analysis of variance (NP-MANOVA) with randomized residuals in a permutation procedure (RRPP; see text for statistical details). d is the distance between means in multivariate space (effect size of difference). Significant differences shown in bold with two (*P* < 0.01) asterisks.

| Color Morph Comparison | $d$ | 95% Upper Confidence Limit | $Z$ | $Pr > d$ |
| --- | --- | --- | --- | --- |
| White: Yellow | 0.994 | 0.812 | 2.476 | **0.008\*\*** |
| White: White-Yellow | 0.882 | 1.187 | 0.558 | 0.285 |
| White-Yellow: Yellow | 0.803 | 1.069 | 0.532 | 0.317 |
| Orange: White | 0.846 | 1.696 | -0.514 | 0.698 |
| Orange: White-Yellow | 1.547 | 1.729 | 1.209 | 0.110 |
| Orange: Yellow | 1.509 | 1.626 | 1.389 | 0.082 |

**Table 2.** Proportion of variance explained and variable loadings of predicted principal component (PC) values for first two axes of variation describing the phenotype of common wall lizards (*Podarcis muralis*) from Ohio, USA. Predicted values were generated using a non-parametric multivariate analysis of variance (NP-MANOVA) with randomized residuals in a permutation procedure (RRPP), including the fixed effects of color morph and sex (see main text for statistical details). Abbreviations: CORT = Plasma corticosterone concentration; TRIG = Plasma triglyceride concentration; Hct = Hematocrit; T_b_ = Field body temperature; SMI = Standardized mass index

|  | PC1 | PC2 |
| --- | --- | --- |
| Proportion variance explained | 75.38% | 20.81% |
| CORT ($\log_{10}$) | -0.631 | -0.728 |
| TRIG ($\log_{10}$) | -0.340 | 0.119 |
| Hct | -0.291 | 0.134 |
| $T_b$ | 0.304 | -0.533 |
| SMI | -0.556 | 0.392 |

**Table 3.** Mean values ± SD of five physiological or behavioral measures common wall lizards (*Podarcis muralis*) from Ohio, USA by color morph.

|  | **CORT**<br>(ng/mL) | **TRIG**<br>(mg/dL) | **Hct** | **T<sub>b</sub>**<br>(°C) | **SMI**<br>(g) |
| --- | --- | --- | --- | --- | --- |
| **White**<br>N = 23 | 11.8 ± 13.1 | 150 ± 79.2 | 0.350 ± 0.0586 | 34.0 ± 3.50 | 5.64 ± 0.550 |
| **White-Yellow**<br>N = 13 | 24.6 ± 36.9 | 154 ± 96.2 | 0.377 ± 0.0846 | 32.5 ± 3.58 | 5.95 ± 0.0894 |
| **Yellow**<br>N = 45 | 40.7 ± 42.0 | 156 ± 93.5 | 0.367 ± 0.0725 | 34.3 ± 2.90 | 5.80 ± 0.697 |
| **Orange**<br>N = 5 | 6.34 ± 3.55 | 111 ± 64.3 | 0.344 ± 0.0735 | 34.6 ± 2.24 | 5.26 ± 0.698 |
| **All**<br>N = 86 | 28.55 ± 36.54 | 151.58 ± 88.11 | 0.362 ± 0.0705 | 33.95 ± 3.16 | 5.75 ± 0.700 |

CORT = Plasma corticosterone concentration; TRIG = Plasma triglyceride concentration; Hct = Hematocrit; T<sub>b</sub> = Field body temperature; SMI = Standardized mass index

The first two axes of variation in the multivariate physiological phenotype varied across seasons. PC1 differed among color morphs, between sexes and between sampling years. Across the season, PC1 shifted linearly in response to day of year in a manner dependent on color morph and shifted in response to the quadratic effect of day of year (Table 4; Fig. 4A). PC2 differed among morphs and between sexes. Across the season, PC2 increased with time of day and increased linearly across the season in a manner consistent across morphs (Table 4; Fig. 4B).

**Figure 4.**
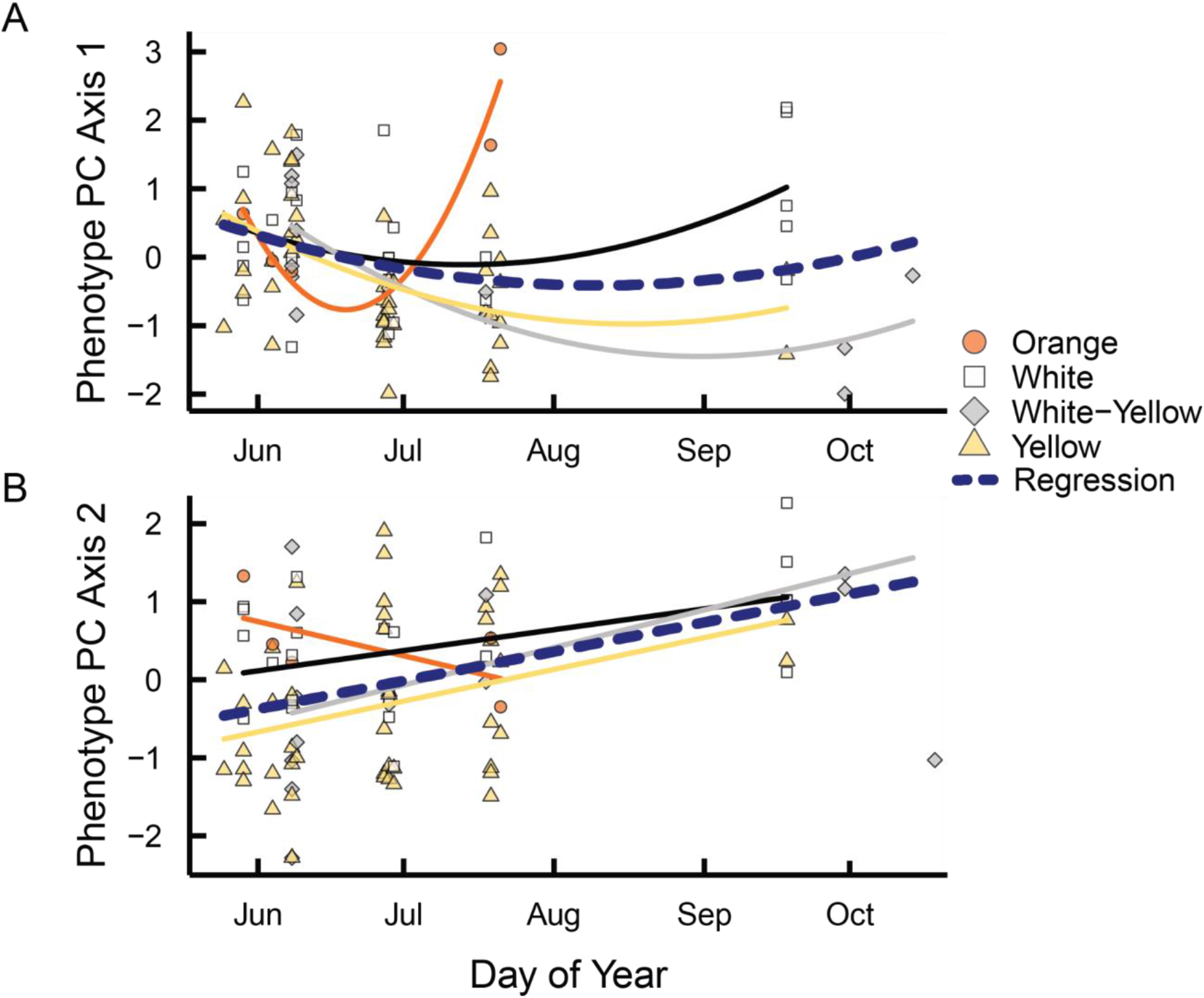
Variation in individual scores on Principal Component Axis 1 (A) and Principal Component Axis 2 (B) across activity season in common wall lizards (*Podarcis muralis*) from Ohio, USA by color morph. Individual PC scores are predicted from non-parametric multivariate analysis of variance (NP-MANOVA) with randomized residuals in a permutation procedure (RRPP), including the fixed effects of color morph and sex (see main text for statistical details). Loadings for each PC axis are provided in Table 2. Regression lines are shown by color morph and for all lizards, with quadratic lines shown in (A) and linear lines in (B), in concordance with linear model results (presented in Table 4).

**Table 4.** Results ocf linear model analysis of the effects of color morph, sex, and time on the first two axes of variation describing the phenotype of common wall lizards (*Podarcis muralis*) from Ohio, USA. Significant differences shown in bold with one (P < 0.05) or two (P < 0.01) asterisks.

| Source of Variation | PC 1 | PC 2 |
| --- | --- | --- |
| Color Morph |  |  |
| Estimate ± SE | White: $5.09 \pm 3.02$<br>White-Yellow: $8.93 \pm 3.22$<br>Yellow: $7.85 \pm 2.93$ | White: $-0.10 \pm 0.41$<br>White-Yellow: $-0.35 \pm 0.41$<br>Yellow: $-0.69 \pm 0.40$ |
| Test statistic: $F(df_n, df_d)$ | 5.41 (3, 77) | 5.08 (3, 81) |
| P-value | **0.0020\*\*** | **0.0028\*\*** |
| Sex |  |  |
| Estimate ± SE | Male: $-0.96 \pm 0.23$ | Male: $0.50 \pm 0.22$ |
| Test statistic: $F(df_n, df_d)$ | 22.04 (1, 77) | 8.33 (1, 81) |
| P-value | **<0.0001\*\*\*** | **0.0050\*\*** |
| Year |  |  |
| Estimate ± SE | $-0.0075 \pm 0.25$ | |
| Test statistic: $F(df_n, df_d)$ | 7.23 (1, 77) | — |
| P-value | **0.0088\*\*** |  |
| Day of Year (linear) |  |  |
| Estimate ± SE | $-0.074 \pm 0.038$ | $0.011 \pm 0.0026$ |
| Test statistic: $F(df_n, df_d)$ | 2.32 (1, 77) | 16.55 (1, 81) |
| P-value | 0.13 | **0.00011\*\*** |
| Day of Year (quadratic) |  |  |
| Estimate ± SE | $0.00028 \pm 0.000097$ | |
| Test statistic: $F(df_n, df_d)$ | 4.73 (1, 77) | — |
| P-value | **0.033\*** |  |
| Time of Day (linear) |  |  |
| Estimate ± SE | | $0.000039 \pm 0.000010$ |
| Test statistic: $F(df_n, df_d)$ | — | 14.05 (1, 81) |
| P-value |  | **0.00033\*\*** |
| Color Morph × Day of Year (linear) | | |
| Estimate ± SE | White: $-0.034 \pm 0.017$<br>White-Yellow: $-0.58 \pm 0.019$<br>Yellow: $-0.51 \pm 0.017$ | |
| Test statistic: $F(df_n, df_d)$ | **7.59 (3, 77)** | — |
| P-value | **0.00016\*\*** |  |

The results of univariate models are consistent with those of the multivariate analyses. Plasma CORT concentration differed among color morphs, with pairwise comparisons indicating significant differences between white and yellow morphs (t_79_ = −3.59, *P* = 0.0031), with a trend for differences between orange and yellow morphs (t_79_ = −2.39, *P* = 0.087). Additionally, TRIG concentration and SMI varied linearly across the active season in a manner dependent on color morph. TRIG values were higher in the 2020 field season, while Hct and SMI were higher in 2021. CORT, Hct, and SMI decreased linearly across the active season; conversely TRIG increased non-linearly across the active season. CORT was higher earlier in the day and decreased linearly across the daily active period. SMI was higher in males compared to females. None of the tested predictors influenced T_b_. All final univariate model results are presented in Table S2.

## Discussion

Our results demonstrate a clear separation in the multivariate physiological phenotype among *P. muralis* color morphs, including traits related to energy processing and storage. As such, these polymorphisms represent distinct physiological and behavioral strategies that continue to coexist in a novel, urban environment after a single transcontinental introduction event. Color polymorphisms, especially in *Podarcis*, represent various combinations of physiological, morphological, and behavioral traits, potentially increasing the breadth of phenotypic options available for species in a new environment and facilitating range expansion (Forsman *et al*. 2008). Color polymorphism has been documented many times in the native range of *P. muralis* (Abalos *et al*. 2020; Abalos *et al*. 2022; Sacchi *et al*. 2013), but little work has been done on the recently-established populations that reside in southern Ohio (Brown *et al*. 1995a; Brown *et al*. 1995b; Vaughn *et al*. 2021). This study is the first to report on the patterns of trait covariation among color morphs in an introduced species. Surprisingly, the initial propagule of only 10 individuals (Deichsel & Gist 2001) must have included the genetic variation necessary to maintain the observed color polymorphisms in their modern-day descendants. This diversity could be responsible for their adaptive success in the novel ecosystems of Cincinnati, as the population has exploded well into the hundreds of thousands (J. Davis pers. comm; Kwiat & Gist 1987), even after undergoing such a severe genetic bottleneck (Homan 2013; Lescano 2010). Other *Podarcis* species that were introduced to the US have undergone similar genetic bottlenecks (Kolbe *et al*. 2013), but despite this, have established thriving populations containing thousands of lizards.

### Phenotypic variation

As with other studies of color polymorphic lizard species (e.g., Calsbeek *et al*. 2010; Galeotti *et al*. 2010; Huyghe *et al*. 2009; Sacchi *et al*. 2007b), we identified a clear separation among color morphs in the physiological-behavioral phenotype (Table 1, Fig. 3). We measured traits specifically relevant as indicators of energetic processing and status at different time scales, as well as thermoregulation which will fundamentally drive the pace of all physiological processes (Angilletta 2009; Black *et al*. 2019). Circulating total triglycerides (TRIG), a measure of near-term energy availability, and standardized mass index (SMI), a measure of long-term energy storage, were positively correlated among individuals (Table 2). These traits were also positively correlated with circulating corticosterone (CORT), contrary to our expectations that CORT would be elevated in low-energy individuals to promote feeding (Gangloff & Greenberg 2023). Our measure of CORT represents a baseline level, because we collected blood samples immediately after capture before CORT becomes elevated due to the stress response (Tylan *et al*. 2020). As such, our results suggest that individuals with high levels of CORT, but still within the allostatic range (McEwen & Wingfield 2003), are optimizing energy acquisition and processing to maintain high levels of available stores, both in the short and long term. Importantly, yellow and white morphs are separated on this axis of variation, such that yellow morphs exhibited higher levels of CORT and TRIG than white, white-yellow, and orange morphs (Fig. 2, Table 3). This result is noteworthy in the context of previous work with *P. muralis* demonstrating that morphs differ in life history strategy such that yellow morphs are r-strategists (produce many eggs with small offspring) in contrast to the K-strategy employed by white morphs (fewer but larger offspring; Galeotti *et al*. 2013). Higher levels of circulating CORT in the faster pace-of-life yellow morphs is contrary to patterns of intraspecific variation in life history and corticosterone in garter snakes (*T. elegans*, Holden *et al*. 2022, Palacios *et al*. 2012), suggesting a lack of broad patterns of covariation among life-history traits and hormonal pathways in squamate reptiles. The second axis of variation is most clearly defined by contrasts between individuals with high CORT and high selected field body temperature (T_b_) and lizards with the opposite pattern. While this axis of variation explains only ∼20% of the variation among color morphs, here we find separation of the white morph from the intermediate white-yellow morph, suggesting that white-yellow morphs represent a strategy intermediate between the ‘pure’ morphs. In addition to this, the intermediate morph was far less common than either the white or yellow morph, suggesting that assortative mating may reduce the frequency of intermediate morphs (Sacchi *et al*. 2018), despite that early-life offspring are equally fit among morph combinations (Abalos *et al*. 2022). Interestingly, this intermediate white-yellow morph exhibited the highest mean SMI, often used as a fitness proxy, in our study (Table 3, Fig. 3).

### Seasonal shifts

In addition to the clear separation of physiological phenotypes between white and yellow morphs, we found morph-specific shifts in physiological traits across the active season (Table 4, Fig. 4). Values of PC1 generally decrease across the active season, though the pattern is non-linear. Early in the season, morphs exhibit similar values on this axis of variation, but by late summer white morphs have higher scores, compared to yellow or white-yellow morphs, indicating a reduction in energy capacity with lower CORT, TRIG, and SMI values. All morphs similarly increased values of PC 2 across the active season (Table 4, Fig. 4), primarily indicating a decrease in CORT and selected body temperature by late summer. It is not surprising that CORT was the most variable trait among morphs, varied strongly across the season, and was the only trait affected by time of day (Table S2). In reptiles, CORT is the primary glucocorticoid in the hypothalamic-pituitary-adrenal axis (reviewed in Gangloff & Greenberg 2023), a hormonal pathway whose fingerprint rests upon a variety of physiological processes related to maintenance of energy balance. As such, hormones of this pathway respond to the perceived environment to maintain organismal homeostasis. Thus, measures of circulating hormones do not represent differences in physiological endpoints so much as differences in strategies to achieve regulation of important physiological parameters, for example circulating glucose (MacDougall-Shackleton *et al*. 2019; Romero & Beattie 2022). Flexibility in these pathways in response to changing or novel conditions is especially relevant in the context of urbanization (Bonier 2023). Nonetheless, we do observe that TRIG and SMI, commonly used as a fitness proxy, both decrease in white morphs by late in the summer. Future work is needed to elucidate the long-term effects of this decrease in available energy, for example in relation to overwinter survivorship or reproduction in the following year.

### Thermoregulation

The fundamental driver of variation in determining the rate of energy processing in all organisms is body temperature (Angilletta 2009). In ectothermic vertebrates which actively thermoregulate, body temperature is maintained through active selection of microsites with favorable temperatures, though of course the ability to attain suitable temperatures is dependent on those available in the environment (Angilletta 2009; Hertz *et al*. 1993). Given the high level of variability in temperate climates, lizards, including *P. muralis*, are often very efficient thermoregulators (Bodensteiner *et al*. 2021b; Ortega *et al*. 2016). Contrary to our prediction that lizard body temperature would vary according to seasonality or time of day, we found that none of our predicted factors influenced T_b_, suggesting that lizards are highly efficient at maintaining their preferred body temperature. The mean field T_b_ we observed here, 34.0°C, is close to the mean active body temperature of lizards from an Italian population in August (33.6°C; Avery 1978) and to the optimal temperature for sprint performance (33.7°C; Telemeco *et al*. 2022) of lizards from populations in France. The slightly higher field body temperatures of Ohio lizards complements other findings that *P. muralis* in Ohio may have altered thermal physiology relative to continental Europe populations, including higher thermal preferences (Spears *et al*. 2023, in review) and an altered thermal tolerance breadth (Litmer & Murray 2019). Surprisingly, we did not find evidence that color morph or sex influenced field body temperatures (Matthews *et al*. 2023). Previous work in the congener *P. erhardii* shows that orange morphs have a dramatically reduced thermal preference compared to other morphs (Thompson *et al*. 2023). Despite sampling across the female reproductive season, which influences thermoregulation in a variety of lizard species (reviewed in: Bodensteiner *et al*. 2021a; Lailvaux 2007), we found no variation in selected body temperatures. Given that our fieldwork was conducted under generally favorable conditions (our goal was to catch lizards when active), it would be enlightening to examine variation in selected body temperatures under suboptimal conditions, for example at night or during overwintering.

### Sexual factors

In other species of lizard, color polymorphism is a factor for sexual dimorphism. In *Podarcis* species however, it is less cut and dry. *P. muralis* color morphs are not relevant to “sociosexual behavior” (Abalos *et al*. 2020), but there is evidence that the different female morphs employ different reproductive strategies (Pellitteri-Rosa 2010). There is no female-male color dimorphism in *Podarcis* species, as both sexes contain the same number and vibrancy of color morphs (Brock *et al*. 2020). In fact, the only documented dimorphism between the sexes is that of body size (Sacchi *et al*. 2015b), though we did not find significant dimorphisms in body size here. Color morph has no impact on sex, and vice versa-female and male color morphs had the same fitness (Abalos *et al*. 2020; Calsbeek *et al*. 2010), the same sprint performance (Zajitschek *et al*. 2012), same use of thermal habitats and thermal preference (although gravid females have lower field body temperature; Braña 1993; Thompson *et al*. 2023; Zajitschek *et al*. 2012), and the same levels of immunocompetence (Calsbeek *et al*. 2010). However, despite this, the natural life history of *P. muralis* includes a reproductive season that affects both the body temperature and the SMI of reproductive females (Braña 1993). We did not find differences between sexes in T_b_, but females exhibited a significantly lower SMI compared to males (Fig. S1, Table S2), most likely due to catching post-oviposition females with depleted energy stores.

### Habitat differences

In interaction with sexual selection, environmental-driven selective pressures may be acting on white, yellow, white-yellow, and orange morphs generating optimum geographic and thermal patterns (Pérez i de Lanuza *et al*. 2018; Zajitschek *et al*. 2012). Morph frequencies have been known to differ spatially, with orange morphs preferring lower temperatures and being more common at high elevations (Pérez i de Lanuza & Carretero 2018). Similar trends have been observed in *P. erhardii* orange morphs, which exhibit lower preferred temperatures and occupy more vegetated microhabitats, suggesting a shared preference for cooler microhabitats in orange morphs across species (BeVier *et al*. 2022; Thompson *et al*. 2023). Further, morphs have been observed to vary in size (Brock *et al*. 2020), with orange morphs being much larger than their white, white-yellow, and yellow counterparts (mean orange SVL: 64.7; mean white, white-yellow, and yellow: 61.6). Though there is a low proportion of orange morphs in Ohio populations of *P. muralis* generally, in the time since these surveys were conducted we have observed a population in Cincinnati with a high proportion of orange morphs at a site with highly manicured vegetation and supplemental watering (*pers. obs.*). Additionally, orange color morphs in introduced populations of *P. siculus* in southern California appear more frequently in shady and irrigated gardens (*pers. obs.*). Future work will be directed toward identifying variation in microhabitat selection among morphs and specifically locating populations with higher orange morph frequencies in areas with supplemental watering and/or vegetation. While our results suggest that *P. muralis* orange morphs may be distinct in a number of aspects of their physiology and ecology, our low sample size (N = 5) precludes definitive conclusions. Nevertheless, these results are consistent with previous observations in other *Podarcis* populations across species that clearly show distinct differences in physiology and behavior in the orange morphs.

## Conclusion

Overall, our results suggest that color morph variation in physiological phenotype can provide several solutions to ecological challenges, and this variation may confer adaptive advantages in small populations introduced to novel environments (Forsman *et al*. 2008). Different strategies associated with color polyphenisms, for example, may allow for increased variation of niche width among individuals within a population, concurrent with increased adaptive potential, without increasing genetic load (sensu Van Valen 1965). Little is known about the adaptive significance of color polymorphism in wall lizards (Abalos *et al*. 2020; Zajitschek *et al*. 2012), despite that the genus *Podarcis* comprises mostly color polymorphic species (Brock *et al*. 2022a). Introduced populations of *Podarcis* provide new opportunities to test longstanding theories and understand the causes and consequences of color polymorphism.

## Acknowledgements

This work was supported by the Ohio Wesleyan University Summer Science Research Program, the Small Grant Program, and a Theory-to-Practice Grant. SS received support from a Roger Conant Grants-in-Herpetology award from the Society for the Study of Reptiles and Amphibians and PLV was supported by a Travel Grant from the Midwestern Partners in Amphibian and Reptile Conservation. KMB was supported by an NSF PRFB (Award number 2109710). We also thank J. Davis, G. Lipps, J. Sockman, G. Hatosky, and Columbus Downtown High School students for assistance with finding lizards and fieldwork, and L. Tabak, J. Arlington, & L. Tuhela-Reuning for logistical support.

---

**Fig. S1.**
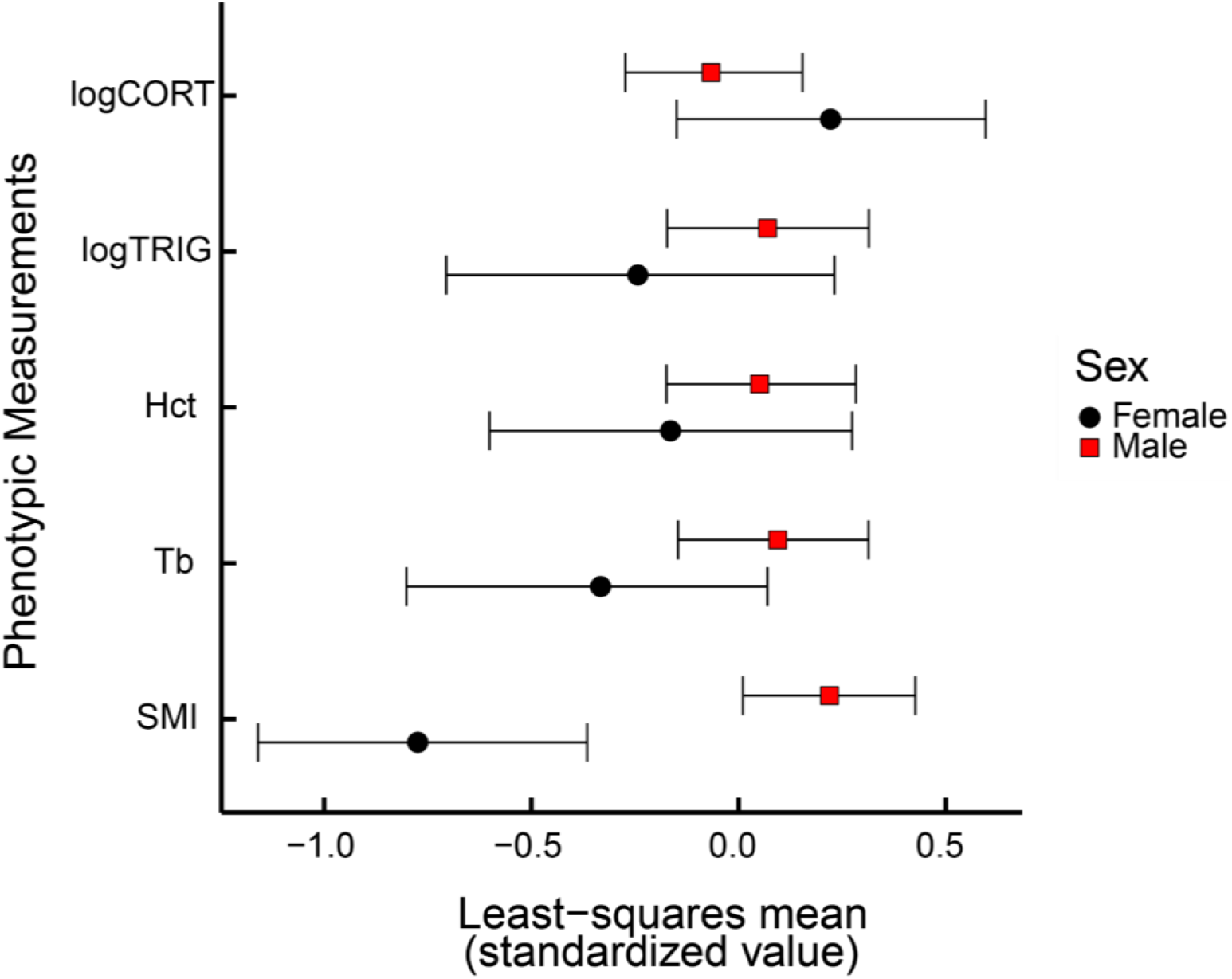
Least-squares means and 95% confidence intervals for phenotypic traits of common wall lizards (*Podarcis muralis*) from Ohio, USA generated from a non-parametric multivariate analysis of variance (NP-MANOVA) with randomized residuals in a permutation procedure (RRPP), including the fixed effects of color morph and sex (see main text for statistical details). Values shown are predicted from the model after accounting for covariation within the response matrix, displayed on a z-standardized scale. Abbreviations: CORT = Plasma corticosterone concentration; TRIG = Plasma triglyceride concentration; Hct = Hematocrit; T_b_ = Field body temperature; SMI = Standardized mass index

**Table S1.** Locality and collection data for common wall lizards (*Podarcis muralis*) from Ohio, USA.

| POPULATION NAME<br>(Abbreviation) | LATITUDE<br>LONGITUDE | ANIMAL<br>COLLECTION<br>DATE | White Morphs<br>(N = 23) | White-Yellow Morphs<br>(N = 13) | Yellow Morphs<br>(N = 45) | Orange Morphs<br>(N = 5) |
| --- | --- | --- | --- | --- | --- | --- |
| ALMS PARK<br>(ALM) | 39°06'41.1"N<br>84°25'49.3"W | 27 June 2021<br>18 July 2021 | N Female = 0<br>N Males = 1 | N Females = 0<br>N Males = 0 | N Females = 1<br>N Males = 2 | N Females = 1<br>N Males = 0 |
| AULT PARK<br>(AUL) | 39°08'06.7"N<br>84°24'24.1"W | 2 June 2020<br>19 July 2020<br>16 September 2020<br>7 June 2021<br>26 June 2021<br>17 July 2021 | N Females = 1<br>N Males = 5 | N Females = 1<br>N Males = 2 | N Females = 1<br>N Males = 9 | N Females = 1<br>N Males = 2 |
| COLUMBUS DOWNTOWN<br>HIGH SCHOOL<br>(CDHS) | 39.9551° N<br>82.9943° W | 29 September 2021<br>13 October 2021 | N Females = 0<br>N Males = 0 | N Females = 0<br>N Males = 3 | N Females = 0<br>N Males = 0 | N Females = 0<br>N Males = 0 |
| MADISONVILLE<br>(MAD) | 39°09'26.0"N<br>84°23'37.1"W | 23 May 2020<br>2 June 2020<br>7 June 2021<br>27 June 2021<br>17 July 2021 | N Females = 0<br>N Males = 3 | N Females = 3<br>N Males = 1 | N Females = 3<br>N Males = 9 | N Females = 0<br>N Males = 0 |
| McMICKEN<br>(MCM) | 39°08'01.7"N<br>84°31'51.0"W | 27 May 2020<br>19 July 2020 | N Females = 0<br>N Males = 0 | N Females = 0<br>N Males = 0 | N Females = 0<br>N Males = 4 | N Females = 0<br>N Males = 0 |
| MISTLETOE<br>(MIS) | 39°06'43.8"N<br>84°33'05.0"W | 27 May 2020<br>2 June 2020<br>8 June 2021<br>28 June 2021<br>18 July 2021 | N Females = 0<br>N Males = 4 | N Females = 0<br>N Males = 1 | N Females = 3<br>N Males = 5 | N Females = 0<br>N Males = 1 |
| RIVER ROAD<br>(RIV) | 39°5' 33.6726"N<br>84°33' 42.9366"W | 27 May 2020<br>16 September 2020<br>8 June 2021<br>26 June 2021 | N Females = 3<br>N Males = 6 | N Females = 1<br>N Males = 1 | N Females = 0<br>N Males = 8 | N Females = 0<br>N Males = 0 |

**Table S2.**
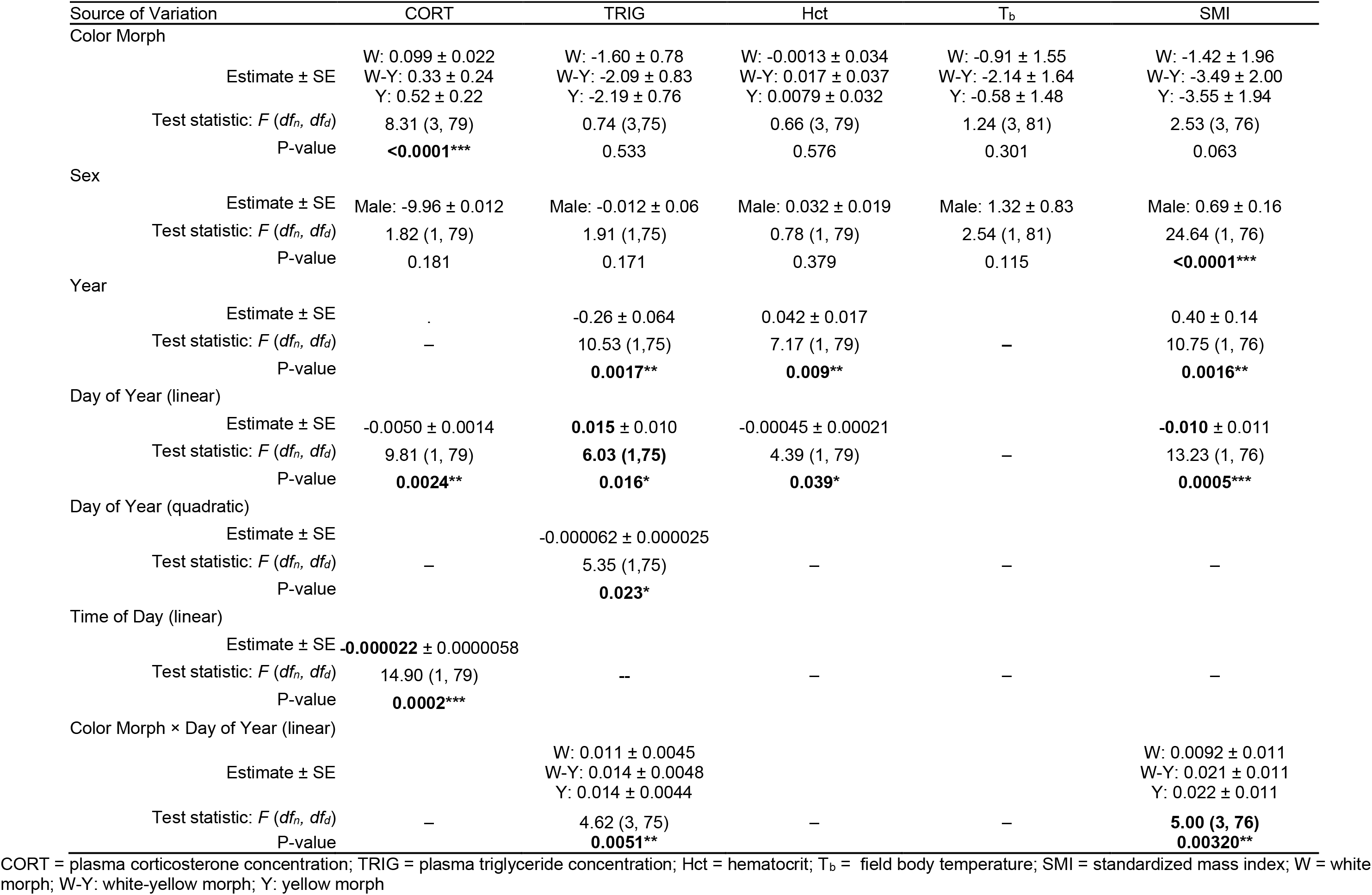
Results of linear model analysis describing the effects of color morph, sex, and time on phenotype measures of common wall lizards (*Podarcis muralis*) from Ohio, USA. Significant differences shown in bold with one (P < 0.05), two (P < 0.01), or three (P < 0.001) asterisks.

## Notes

### Competing Interest Statement

The authors have declared no competing interest.

